# An assessment of the interactions between climatic conditions and genetic characteristic on the agricultural performance of soybeans grown in Northeast Asia

**DOI:** 10.1101/425801

**Authors:** Myoung Ryoul Park, Chunmei Cai, Min-Jung Seo, Hong-Tae Yun, Soo-Kwon Park, Man-Soo Choi, Chang-Hwan Park, Jung Kyung Moon

## Abstract

Glycine max, commonly known as soybean or soya bean, is a species of legume native to East Asia. The interactions between climatic conditions and genetic characteristic affect the agricultural performance of soybean. Therefore, an investigation to identify the main elements affecting the agricultural performances of 11 soybeans was conducted in Northeast Asia, China [Harbin (45°12′N) Yanji (42°53′N) Dalian (39°30′N) Qingdao (36°26′N)] Republic of Korea [Suwon (37°16′N) and Jeonju (35°49′N)]. The days to flowering (DTF) of soybeans with the e1-nf and e1-as alleles and the E1e2e3e4 genotype, except Keumgangkong, Tawonkong, and Duyoukong, was relatively short compared to soybeans with other alleles. Although DTF of the soybeans was highly correlated to all climatic conditions, days to maturity (DTM) and 100-seed weight (HSW) of the soybeans showed no significant correlation with any climatic conditions. The soybeans with a dominant Dt1 allele, except Tawonkong, had the longest stem length (STL). Moreover, the STL of the soybeans grown at the test fields showed a positive correlation with only day length (DL) although the results of our chamber test showed that STL of soybean was positively affected by average temperature (AVT) and DL. Soybean yield (YLD) showed positive correlations with latitude and DL (except L62-667, OT89-5, and OT89-6) although the response of YLD to the climatic conditions was cultivar-specific. Our results show that DTF and STL of soybeans grown in Northeast Asia are highly affected by DL although AVT and genetic characteristic also affect DTF and STL. Along with these results, we confirmed that the DTM, HSW, and YLD of the soybeans vary in relation to their genetic characteristic.

## Introduction

Variability in climate conditions is a major fluctuating factor for estimating crop development. Although the influence of climate on crops depends on geographic location and production conditions [1], interactions between genetic characteristic and climatic conditions, including temperature and precipitation, also affect the agricultural performance of crops [2]. Climatic conditions are consistently changing in Northeast Asia, which is well known as a native geographic range for soybean production. Agronomic scientists and farmers have had to change their crop management practices and adapt their crops to the changing conditions. As is evident from former studies, crop genetics that interact with climate changes have changed due to breeding efforts and genetic engineering.

Numerous researchers have confirmed that climate is also closely related to the agricultural performance of soybean (Glycine max (L.) Merr.) [1, 3, 4]. Soybean is a short-day plant and requires a certain day length (DL) or shorter during its developmental time, including flowering [5, 6]. Despite similarities among soybeans, it has been observed that flowering time varies with environmental conditions [4, 7]. For the first time, Garner and Allard [5] proposed an interactive effect of DL and temperature on soybean floral development, indicating that flowering of soybean is controlled by both photoperiod and temperature. In addition, water deficiency in soybean fields reduces the efficiency of photosynthesis [8], leaf area, crop growth rate, shoot dry matter [9], number of pods, seeds per pod [10], number of flowers during the flowering period, and seed weight during the grain filling stage [11]. Of the yield components, the number of pods is the most sensitive to drought stress and seed weight is least affected [12]. A previous study reported that increased temperature resulted in increased plant height and number of nodes of soybeans [13].

However, along with climatic conditions, genotypic performance generally controls the potential and risk associated with crop production. Genotyping is one of the processes used to confirm differences in the genotypic performances of an individual plant. It has been reported that days to flowering (DTF) and days to maturity (DTM) in soybean are controlled by seven major loci with two alleles at each locus: E1 and E2 [14], E3 [15], E4 [16], E5 [17], E6 [18], E7 [19], E8 [20], and a determinate growth habit locus (Dt1) [21]. Major genes and quantitative trait loci (QTLs) are closely related to each other in the flowering interaction that controls the time to flowering [22]. Consequently, DTF is influenced by a combination of genes and QTLs. Previous studies demonstrated that the determinate stem (Dt1) locus plays an essential role in soybean determination. Stem termination has also been observed to be significantly closed to stem length and maturity of soybean [23], indicating that plant height and maturity might share a similar genetic locus [24, 25]. To examine the genetic characteristic of soybean, many researchers have identified major genes and QTLs controlling agronomic traits including maturity [26] and plant height [27] using map-based cloning. Simple sequence repeat (SSR), an effective genetic marker with various applications, is one of the most favored molecular markers for plant genotyping due to its high polymorphism [28]. Genetic diversity and the genotype of soybeans were also recently analyzed using SSR markers [29, 30, 31].

Several experimental field studies have been performed to analyze the interactions between climatic conditions and genetic characteristic of soybean. In addition, greenhouse and growth chamber tests were conducted to investigate the responses of soybeans to the timing and severity of climatic conditions (e.g., water and temperature stresses), and to complement the results of the field studies [32, 33]. These investigations were geared towards understanding the characteristic of soybeans by examining the effects of climatic conditions and/or genetic characteristic on agricultural performance. However, the interactions between genetic characteristic of soybean and climatic conditions require further assessment. Therefore, the goal of this study was to evaluate the interactions between climatic conditions and genetic characteristic, to inform soybean management and development of new varieties that can adapt to the climate changes in Northeast Asia. We conducted experimental field tests in six locations in Northeast Asia with diverse climatic conditions, growth chamber tests under six combinations of temperature and DL, and a genetic tree analysis constructed using SSR markers that are significantly associated with each agronomic trait of soybeans. These methods allowed a more comprehensive conclusion to be drawn.

## Materials and methods

### Experimental field plot and growth chamber tests

We conducted preliminary tests in 2014 and 2015 to select appropriate soybeans and experimental test fields. We selected 11 varieties with diverse DTF (Hannamkong, Keumgangkong, Tawonkong, Duyoukong, L62-667, OT93-28, OT89-5, OT94-37, OT89-6, OT94-39, and OT94-51) and 6 appropriate experimental fields in China [Harbin (45°12′N, 127°19′E), Yanji (42°53′N, 129°29′E), Dalian (39°30′N, 122°10′E) and Qingdao (36°26′N, 120°03′E)] and the Republic of Korea [Suwon (37°16′N, 127°01′E) and Jeonju (35°49′N, 127°09′E)] (Fig 1). The experimental plots were completely randomized for this study. In 2016 and 2017, we sowed seeds of the soybeans in the middle of May and harvested in the middle of October for the northern sites (Harbin, Yanji and Dalian), and sowed in the middle of June and harvested in the early of October for the southern sites (Suwon, Qingdao, and Jeonju). The seeds were sown in a 70×15 cm spacing in the plots of the six experimental fields and were treated with a basic granular fertilizer of N-P_2_O_5_-K_2_O=40-70-60 kg/ha before sowing.

**Fig 1.**
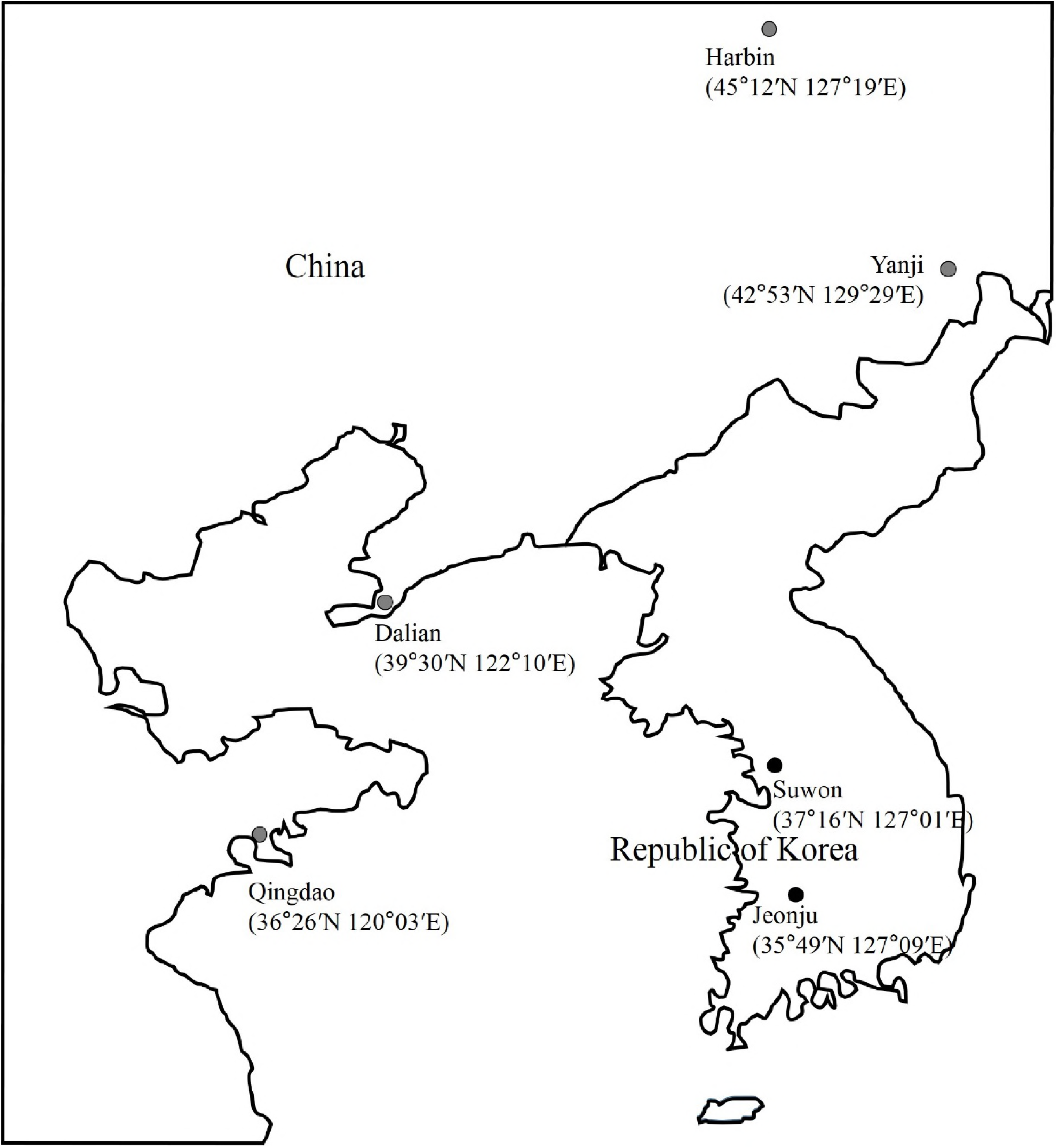
Geographic locations of six experimental sites in China and the Republic of Korea.

To complement the results of the field tests, the DTF and stem length (STL) of soybeans grown in walk-in chambers (Koito manufacturing Co., Ltd., Tokyo, Japan) were investigated. We selected four soybeans based on their DTF tested in the field: Hannamkong, Tawonkong, OT93-28, and OT89-5. The chambers were controlled with six combinations of temperatures (18, 23, and 28°C) and DLs (10 and 15 h) under a fixed relative humidity of 65% from sowing to harvesting of the soybeans.

### Investigation of agricultural performance and climatic condition

Agronomic traits related to soybean growth and yield were measured according to the agricultural science technology standards for investigation of research of the Rural Development Administration (RDA), Republic of Korea [34]. These include: 1). days to flowering (DTF) defined as the days to stage (R1) of the beginning bloom on the main stem after sowing; 2). days to maturity (DTM) defined as days from R1 to stage (R8) of fully maturity (95% of the pods reaching the mature pod color); 3). 100-seed weight (HSW) defined as the weight of 100 seeds measured under 13% moisture content; 4). stem length (STL) defined as the length from the cotyledonary node to the top of the main stem; and 5). yield (YLD, kg/10 a) calculated as the total weight (kg) of seeds harvested in 1 m^2^ × 1000.

Regional climatic conditions at the test fields during the overall growth periods of the soybeans, including DL, average temperature (AVT), accumulated temperature (ACCT), and precipitation (PRC), were investigated using weather data obtained from the Chinese Meteorological Administration and the Korean Meteorological Administration. We examined the data for each climatic element by month then classified the data largely into two soybean growth periods: vegetative (early) and reproductive (late) periods. This classification is because the soybean requires different cultural environmental conditions for these developmental stages [35]. The early period from May to July is commonly the vegetative stage of the soybeans, while the late period from August to October is, on average, the reproductive stage of the soybeans grown in the six test fields.

### Analysis of genetic relationships

We conducted genotyping of the soybeans to estimate the correlation between genetic characteristic and agricultural performances. Genotyping of the soybeans occurred using the allele-specific DNA markers (Table 1) that distinguish recessive alleles from those that are dominant at the maturity loci (E1, E2, E3 and E4), and a determinate growth habit locus (Dt1) [36]. Additionally, we constructed an Unweighted Pair Group Method with Arithmetic Mean (UPGMA) genetic tree using SSR markers that are significantly associated with each trait (Table 2; [37, 38, 39]). The tree was visualized using MEGA version 4.0 [40].

**Table 1.**
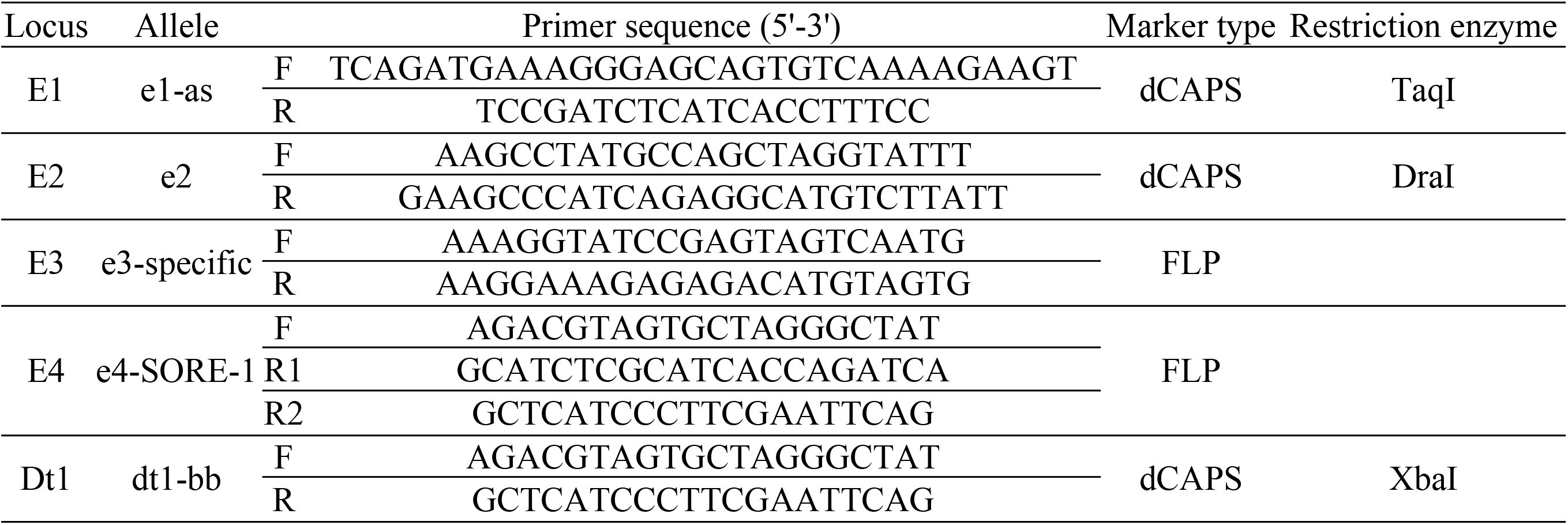
Allele-specific DNA markers that distinguish alleles at the maturity loci (E1, E2, E3 and E4), and a determinate growth habit locus (Dt1) in soybean

**Table 2.** SSR markers associated with days to flowering (DTF), days to maturity (DTM), stem length (STL), 100-seed weight (HSW), and yield (YLD) for the analysis of the genetic relationship.

| Trait | Marker | LG | No. of Chr | Position(cM) | Repeat | Reference |
| --- | --- | --- | --- | --- | --- | --- |
| DTF | Satt322 | C2 | Chr.06 | 82.23 | (ATT)10 | Hu et al., 2014 |
|  | Satt636 | M | Chr.07 | 5 | (ATT)22 | Hu et al., 2014 |
|  | Satt150 | M | Chr.07 | 18.58 | (ATT)20 | Hu et al., 2014 |
|  | Satt417 | K | Chr.09 | 46.2 | (ATT)18 | Hu et al., 2014 |
|  | Satt405 | J | Chr.16 | 12.41 | (ATT)9 | Hu et al., 2014 |
|  | satt285 | J | Chr.16 | 25.51 | (ATT)19 | Hu et al., 2014 |
|  | satt564 | G | Chr.18 | 57.32 | (ATT)22 | Hu et al., 2014 |
|  | Satt594 | G | Chr.18 | 52.94 | (ATT)23 | Rodrigues et al., 2016 |
| DTM | Satt408 | D1a | Chr.01 | 106.69 | (ATT)19 | Hu et al., 2014 |
|  | Satt521 | N | Chr.03 | 65.46 | (ATT)12 | Hu et al., 2014 |
|  | sat_304 | N | Chr.03 | 77.1 | (AT)31 | Hu et al., 2014 |
|  | satt718 | C1 | Chr.04 | 73.79 | (ATC)15, (ATT)15 | Hu et al., 2014 |
|  | sctt010 | G | Chr.18 | 45.87 | (CTT)8 | Hu et al., 2014 |
|  | Satt191 | G | Chr.18 | 96.57 | (ATT)18 | Rodrigues et al., 2016 |
|  | satt614 | I | Chr.20 | 31.94 | (ATT)37 | Hu et al., 2014 |
| STL | satt342 | D1a | Chr.01 | 48.14 | (ATT)21 | Hu et al., 2014 |
|  | Satt436 | D1a | Chr.01 | 70.69 | (ATT)28 | Rodrigues et al., 2016 |
|  | satt641 | N | Chr.03 | 29.28 | (ATT)10 | Hu et al., 2014 |
|  | sat_137 | A1 | Chr.05 | 0 | (AT)16 | Hu et al., 2014 |
|  | satt385 | A1 | Chr.05 | 64.74 | (ATT)22 | Hu et al., 2014 |
|  | Satt454 | A1 | Chr.05 | 28.08 | (ATT)17 | Rodrigues et al., 2016 |
|  | satt389 | D2 | Chr.17 | 79.23 | (ATT)22 | Hu et al., 2014 |
|  | Satt191 | G | Chr.18 | 96.57 | (ATT)18 | Rodrigues et al., 2016 |
|  | Satt354 | I | Chr.20 | 46.22 | (ATT)14 | Rodrigues et al., 2016 |
| HSW | Satt370 | D1a | Chr.01 | 60.99 | (ATT)13 | Rodrigues et al., 2016 |
|  | satt227 | C2 | Chr.06 | 26.65 | (ATT)9 | Hu et al., 2014 |
|  | satt210 | M | Chr.07 | 112.08 | (ATT)25 | Hu et al., 2014 |
|  | Satt536 | M | Chr.07 | 62.14 | (ATT)20 | Rodrigues et al., 2016 |
|  | sat_334 | H | Chr.12 | 59.01 | (AT)21 | Hu et al., 2014 |
|  | satt516 | F | Chr.13 | 44.42 | (ATT)19 | Hu et al., 2014 |
|  | satt706 | E | Chr.15 | 43.36 | (ATT)23 | Hu et al., 2014 |
|  | Satt263 | E | Chr.15 | 45.4 | (ATT)19 | Rodrigues et al., 2016 |
|  | Satt529 | J | Chr.16 | 41.9 | (ATT)13 | Rodrigues et al., 2016 |
|  | Satt154 | D2 | Chr.17 | 57.07 | (ATT)20 | Rodrigues et al., 2016 |
|  | Satt303 | G | Chr.18 | 53.42 | (ATT)20 | Rodrigues et al., 2016 |
|  | Satt562 | I | Chr.20 | 22.84 | (ATT)18 | Rodrigues et al., 2016 |
|  | Satt419 | I | Chr.20 | 21.9 | (ATT)22 | Rodrigues et al., 2016 |
| YLD | Satt216 | D1b | Chr.02 | 9.8 | (ATT)19 | Sebastian et al. 2010 |
|  | satt641 | N | Chr.03 | 29.28 | (ATT)10 | Hu et al., 2014 |
|  | Satt521 | N | Chr.03 | 65.46 | (ATT)12 | Hu et al., 2014 |
|  | Satt190 | C1 | Chr.04 | 73.32 | (ATT)15 | Sebastian et al. 2010 |
|  | satt316 | C2 | Chr.06 | 127.67 | (ATT)12 | Hu et al., 2014 |
|  | Satt307 | C2 | Chr.06 | 121.27 | (ATT)12 | Sebastian et al. 2010 |
|  | Satt429 | A2 | Chr.08 | 162.03 | (ATT)25 | Sebastian et al. 2010 |
|  | Satt556 | B2 | Chr.14 | 73.21 | (ATT)14 | Sebastian et al. 2010 |
|  | Satt594 | G | Chr.18 | 52.94 | (ATT)23 | Sebastian et al. 2010 |
|  | Satt352 | G | Chr.18 | 50.53 | (ATT)18 | Sebastian et al. 2010 |
|  | sct_010 | L | Chr.19 | 59.52 | (CT)10 | Hu et al., 2014 |
|  | Satt497 | L | Chr.19 | 33.71 | (ATT)22 | Sebastian et al. 2010 |
|  | Satt292 | I | Chr.20 | 82.78 | (ATT)16 | Sebastian et al. 2010 |

### Statistical analysis

The Pearson Correlation Coefficient (PCC; Prob>|r| under H_0_: Rho=0, p>0.05) between agricultural performance of the soybean and/or climatic conditions were analyzed with SAS 9.2 (SAS Institute Inc., Cary, NC, USA) and used to assess the degree of correlation. Multiple comparisons of the soybeans within the experimental test fields were performed using the least significant difference (LSD; p>0.05; total degrees of freedom (DF)=30 or 24).

## Results

### Correlation between climatic conditions

DL of the test fields showed a highly positive correlation with latitudes of the fields; a higher latitude results in a longer DL (Fig 2, and S1 Table). Differences in late DL between the test fields were very small and differences in ACCT between the fields were also low (Table 3). AVT showed a negative correlation with the latitudes. PRC and AVT also were negatively correlated with DL (Fig 2, and S1 Table). In Particular, latitude and DL showed highly positive correlations with late PRC and late ACCT but no correlations with PRC and ACCT (Fig 2, and S1 Table). PRC in Suwon (37°16′N) was the highest among the regions and Harbin showed the lowest PRC. AVT was the highest in Qingdao (36°26′N) and Dalian (39°30′N) had the highest ACCT (Fig 2).

**Fig 2.**
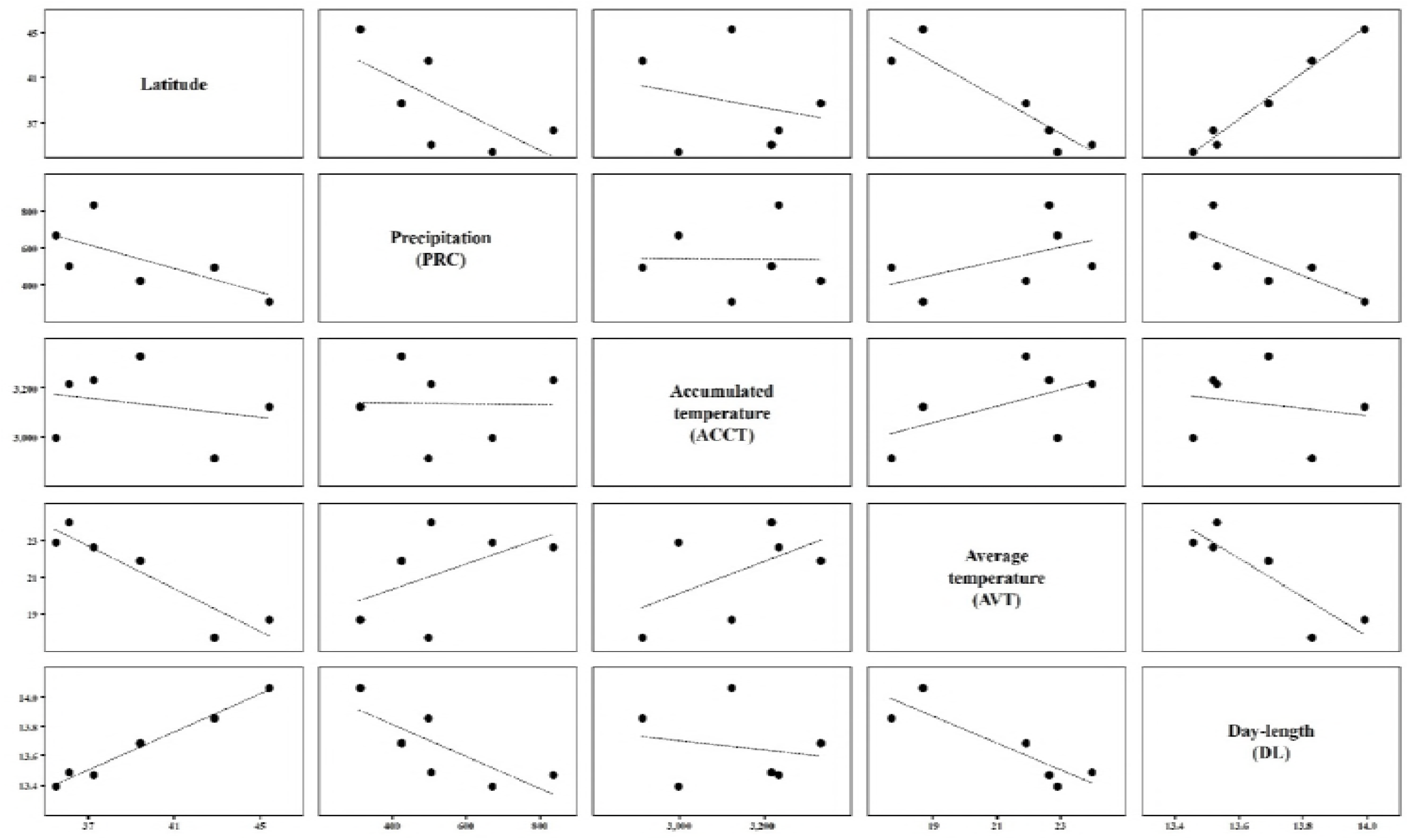
Linear regression analysis between climatic conditions of six experimental sites in China and the Republic of Korea.

**Table 3.**
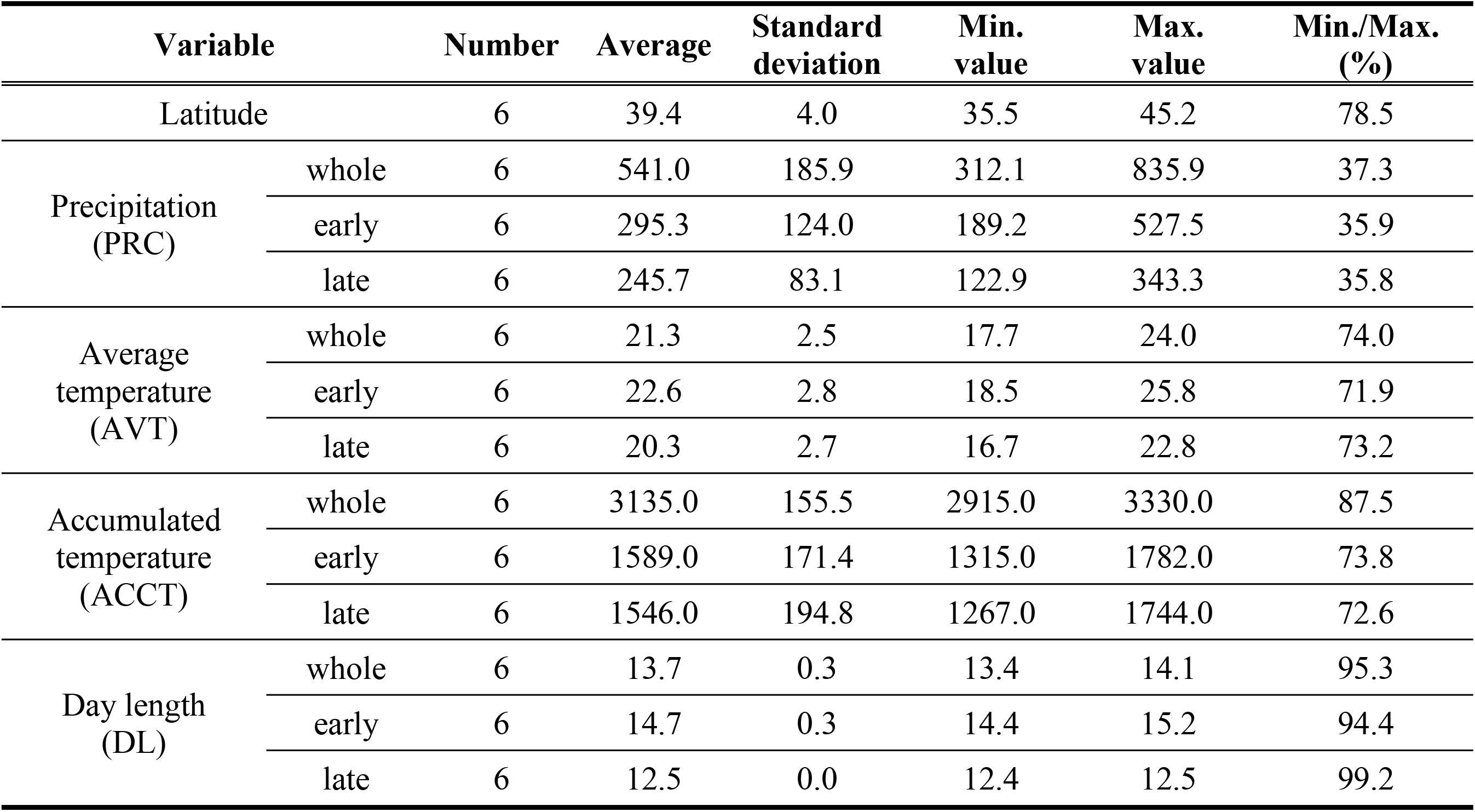
Statistical data of latitude, precipitation (PRC), average temperature (AVT), accumulated temperature (ACCT), day length (DL), and yield (YLD) for six experimental sites in China and the Republic of Korea.

### Correlations between agricultural performances

Pods of four cultivars grown at Harbin (Hannamkong, Keumgangkong, Tawonkong, and Duyoukong) were matured after the frost-free period and, therefore, we could not measure the DTM, HSW, and YLD for these soybeans. DTF of all the soybeans showed positive correlations with DTM, STL, and YLD; however, YLD was negatively correlated with HSW (Fig 3, and S2 Table). Regional differences in HSW of the soybeans were mostly very low compared to the other traits; however, differences between DTF values were the largest (Table 4). The DTF of Hannamkong was the longest among the soybeans while OT89-5 had the shortest DTF and the difference in DTF between them was 22.9 days (Table 4). The DTM of Hannamkong was the shortest although it showed the longest DTF, while OT94-39 had the longest DTM (69.4 days). OT93-28 had the tallest STL; OT94-37 was the shortest length. In the case of YLD, Tawonkong had the highest values and OT93-28 had the lowest (Table 4).

**Fig 3.**
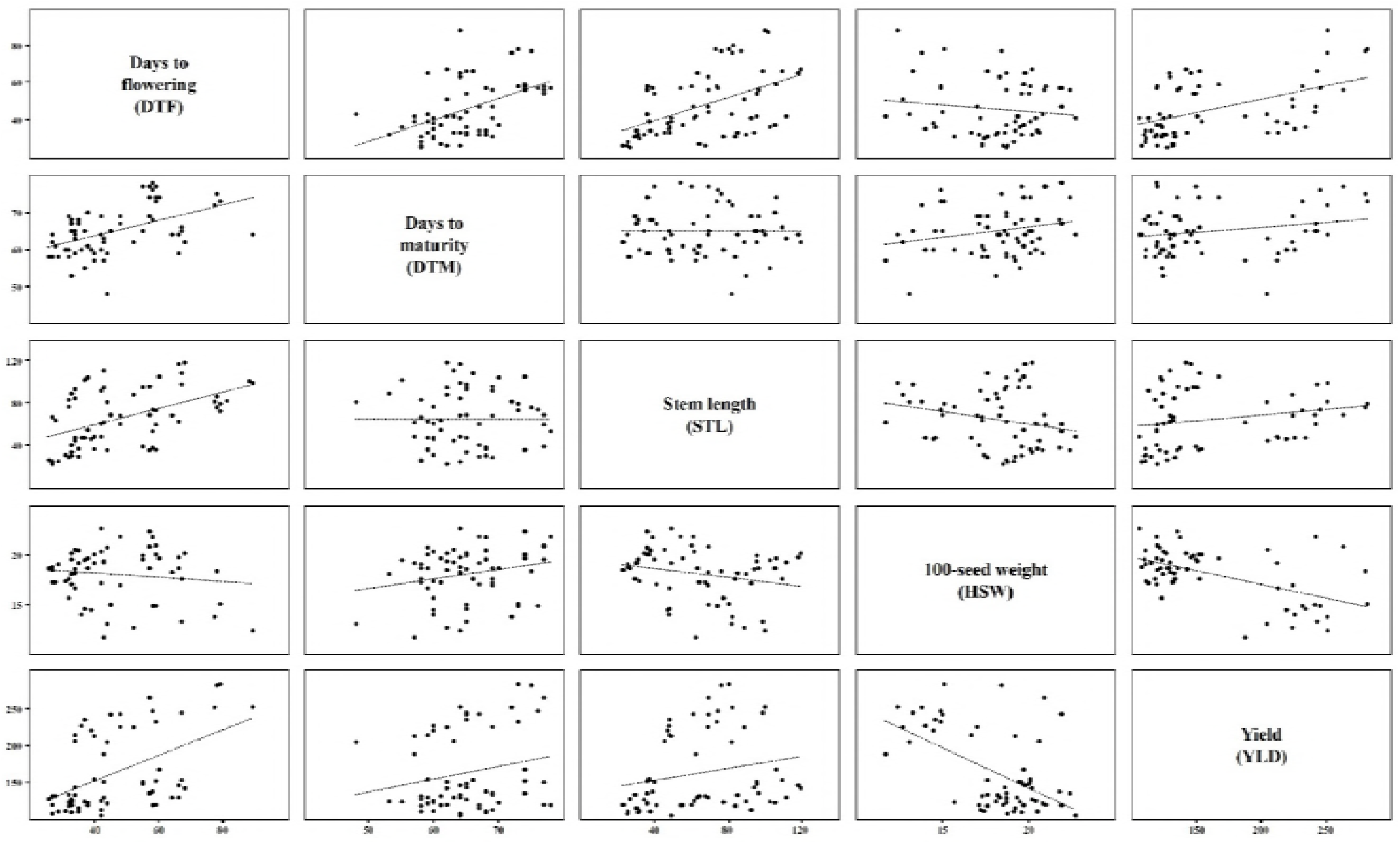
Linear regression analysis between agricultural performances of soybeans grown at six experimental fields in China and the Republic of Korea.

**Table 4.**
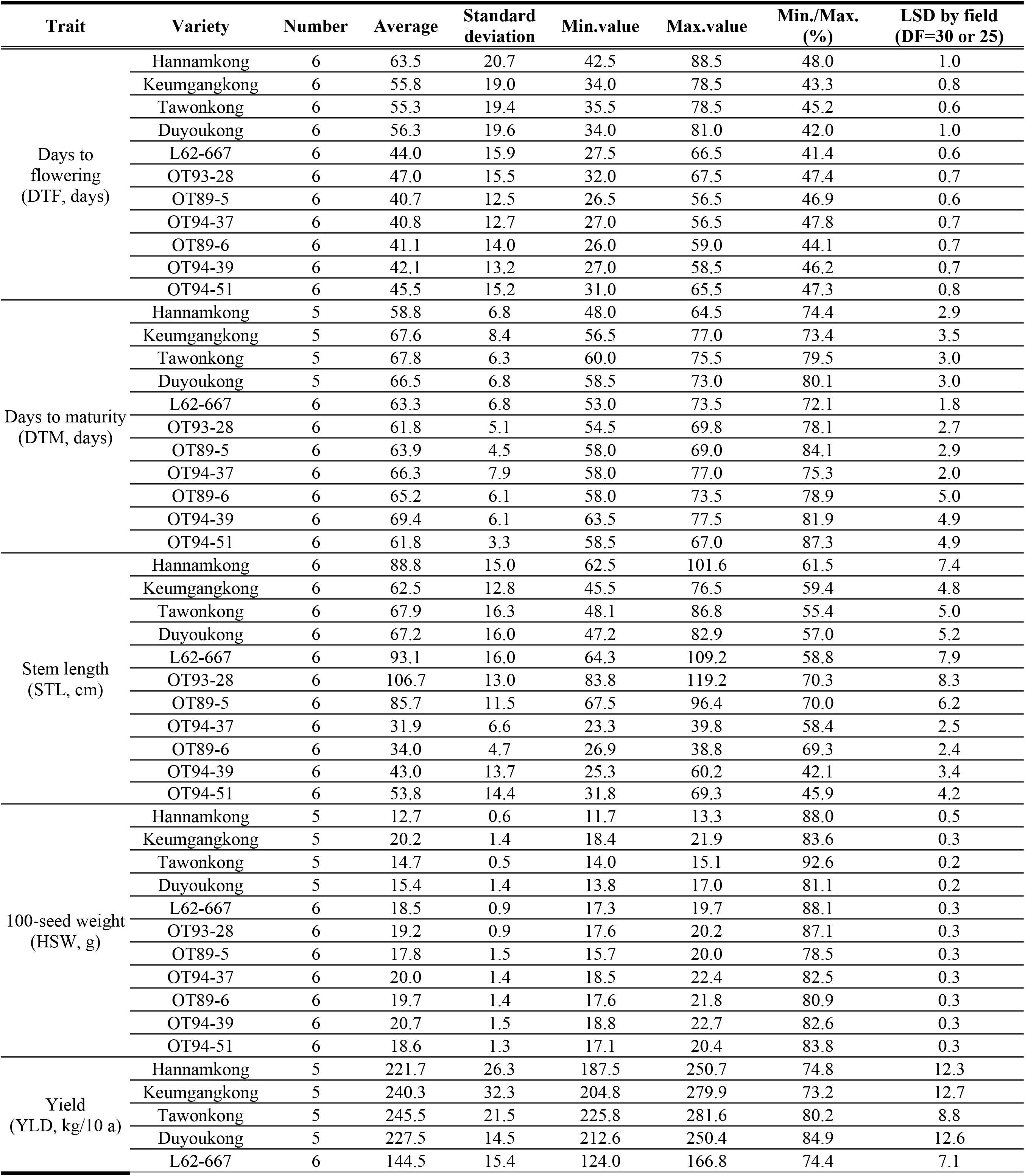
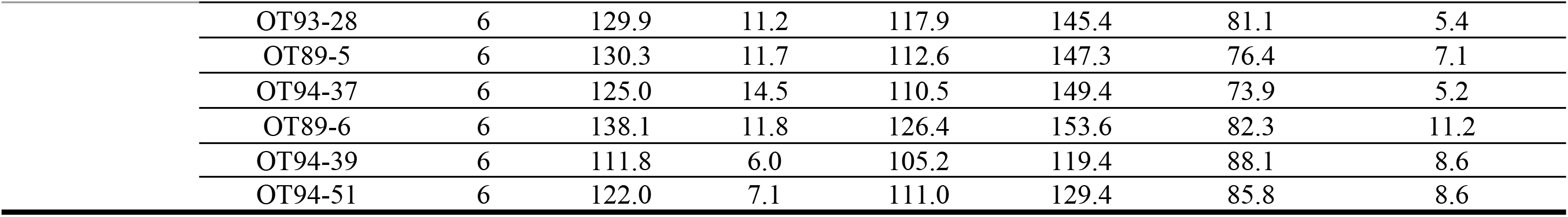
Statistic data for days to flowering (DTF), days to maturity (DTM), stem length (STL), 100-seed weight (HSW), and yield (YLD) of the soybeans grown at six experimental sites in the China and the Republic of Korea.

### Genetic relationships and genotypes

The genetic trees of the soybeans were based on SSR markers that are significantly associated with the agronomic traits. These trees were differentially constructed by traits (Fig 4). The tree based on SSR markers related to DTF had 5 subclades at a genetic distance of 0.5 (Fig 4A). The subclade I, the subclade that Hannamkong is a part of, was very different genetically from the other subclades (Fig 4A). Soybeans in subclades II and III showed the shortest DTF (Fig 4A and Table 4). Genotypes performed using four major E genes (E1 to E4) and stem growth habit locus (Dt1) were significantly different between each soybean (Table 5). The DTF of soybeans that had predominantly E3 and E4 alleles was more than 55 days; Tawonkong only had the E4 allele and had a DTF of 55.3 days (Tables 4 and 5). The tree associated with DTM also formed five subclades at a genetic distance of 0.5 (Fig 4B). Hannamkong, which had the longest DTF and the shortest DTM, was categorized in the subclade III in the tree related to DTM (Table 4 and Fig 4B). The tree associated with STL showed six subclades at a genetic distance of 0.5 (Fig 4C). In the tree associated with STL, the subclade VI (including L62-667, OT89-5, and OT93-28) had the longest STL among the subclades on average; however, the subclade V (including OT94-37 and OT89-6) showed the shortest STL (Fig 4C and Table 4). The soybeans that had a recessive dt1 allele, except Keumgangkong and Duyoukong, had STL of less than 55 cm (Tables 4 and 5). The tree associated with HSW formed three subclades and subclade I, including Hannamkong, Tawonkong, and Duyoukong, was located furthest, genetically, from the other subclades (Fig 4D). The tree associated with YLD also showed three subclades and subclade I, including Hannamkong and Keumgangkong, differed genetically from the other subclades (Fig 4E).

**Fig 4.**
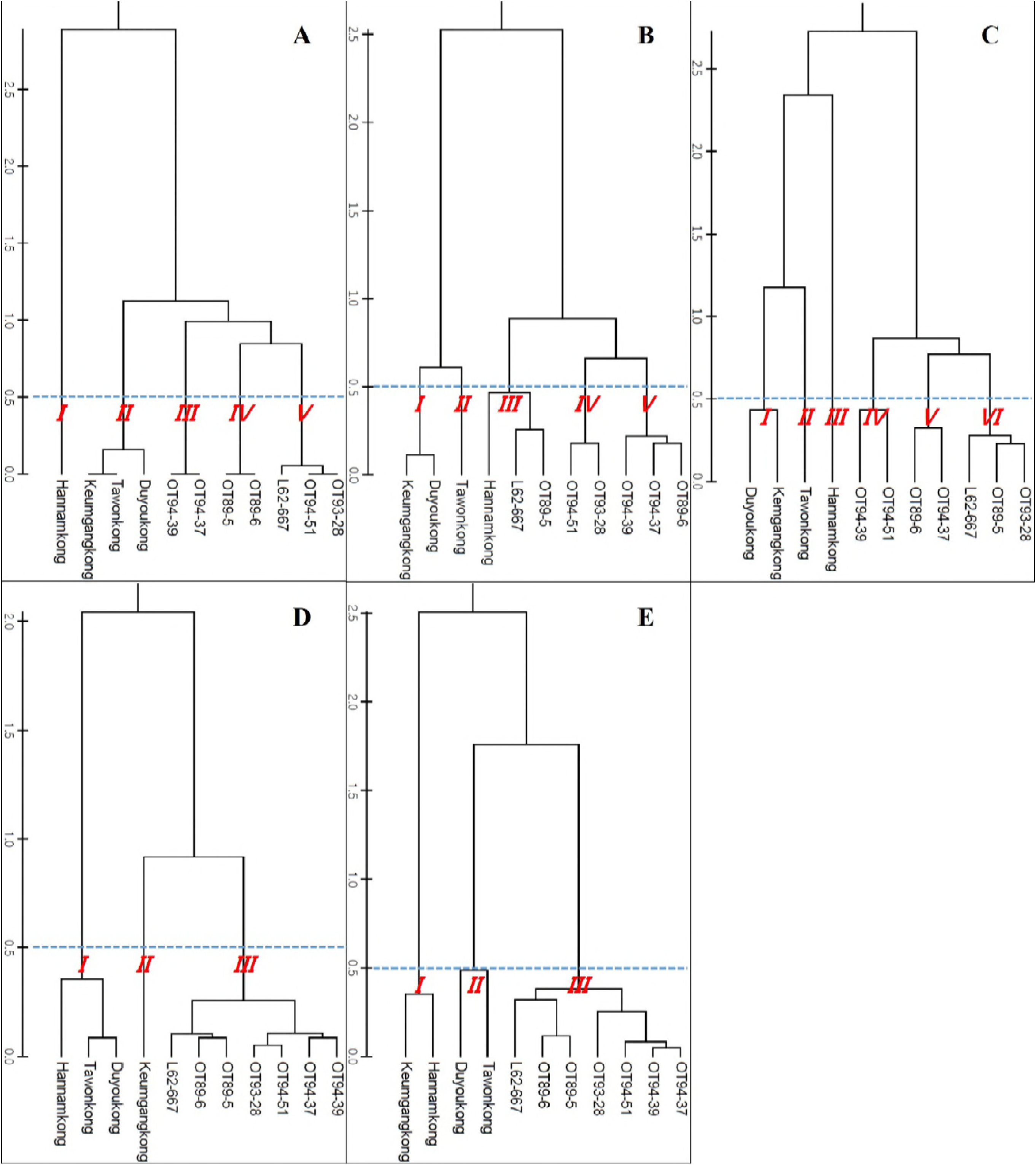
Genetic trees of the soybeans based on SSR markers significantly associated with agronomic traits [A: days to flowering (DTF), B: days to maturity (DTM), C: stem length (STL), D: 100-seed weight (HSW), and E: Yield (YLD)]

**Table 5.** Genotypes of the soybeans grown at six experimental sites in China and the Republic of Korea.

| Variety | Genotype |
| --- | --- |
| Hannamkong | *E1e2E3E4Dt1* |
| Keumgangkong | *e1E2E3E4dt1* |
| Tawonkong | *e1e2e3E4Dt1* |
| Duyoukong | *e1E2E3E4dt1* |
| L62-667 | *e1e2e3E4Dt1* |
| OT93-28 | *E1e2e3e4Dt1* |
| OT89-5 | *e1e2e3e4Dt1* |
| OT94-37 | *e1e2e3E4dt1* |
| OT89-6 | *e1e2e3e4dt1* |
| OT94-39 | *e1e2E3e4dt1* |
| OT94-51 | *E1e2e3e4dt1* |

### Correlations between climatic conditions and agricultural performances

DTF of the soybeans showed high negative and positive correlations with AVT and DL, respectively. DTF showed no significant correlations with PRC or ACCT (Fig 5, and Table 6). STL showed positive correlations with latitude and DL; however, a negative correlation with late PRC was observed (Fig 6, and Table 6). DTM and HSW showed no significant correlation with any of the climatic conditions (Fig 7 and 8, and Table 6). YLD also showed positive correlations with latitude and DL, except in L62-667, OT89-5, and OT89-6 (Fig 9 and Table 6). The soybeans were largely separated into two groups by yield without reference to climatic conditions: the first group included Hannamkong, Keumgangkong, Tawonkong, and Duyoukong, while the second included L62-667, OT93-28, OT89-5, OT94-37, OT89-6, OT94-39, and OT94-51 (Fig 9).

**Fig 5.**
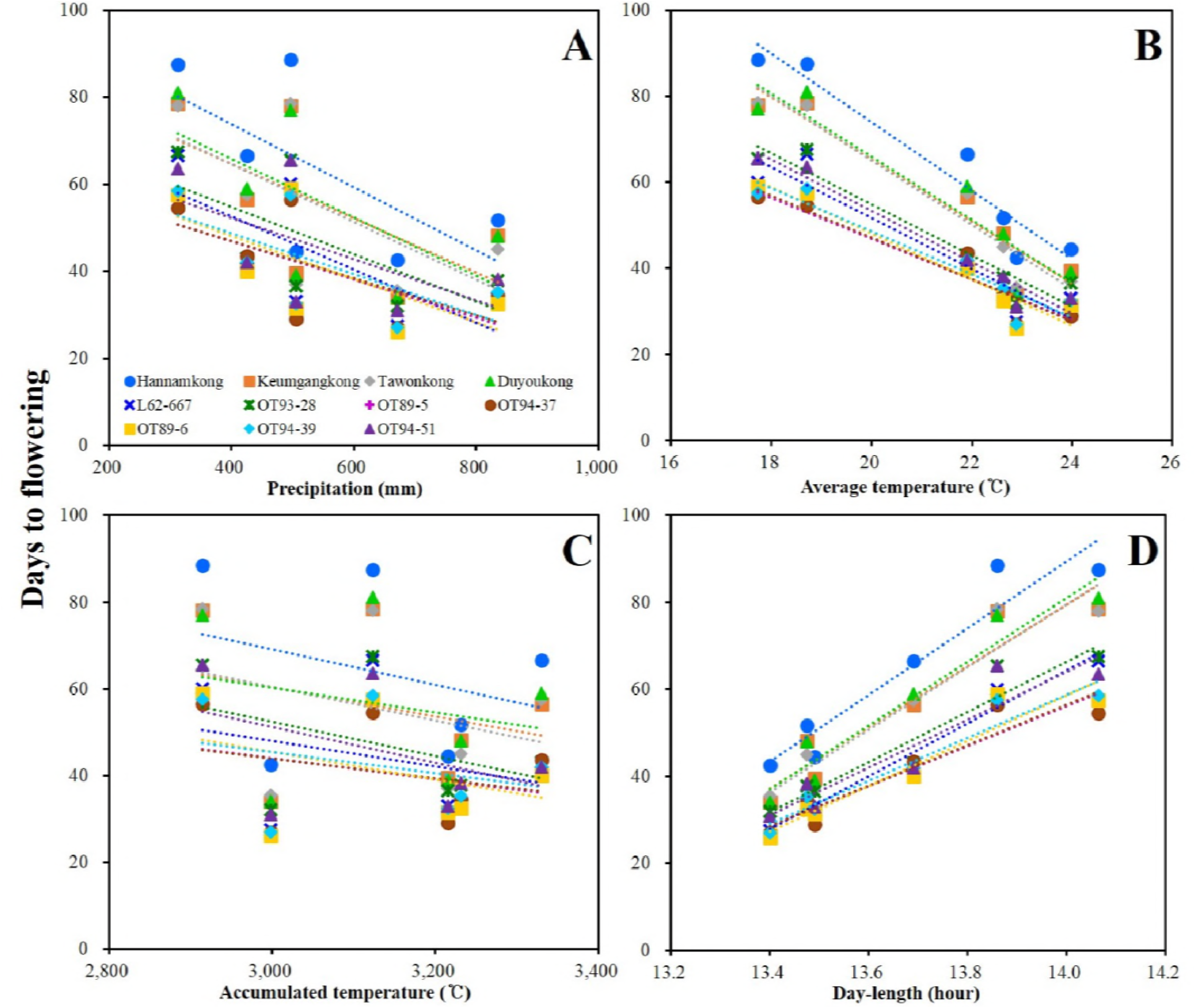
Linear regression analysis between climatic conditions (A: precipitation, B: average temperature, C: accumulated temperature, D: day length) and days to flowering (DTF) of the soybeans grown at six experimental fields in China and the Republic of Korea.

**Fig 6.**
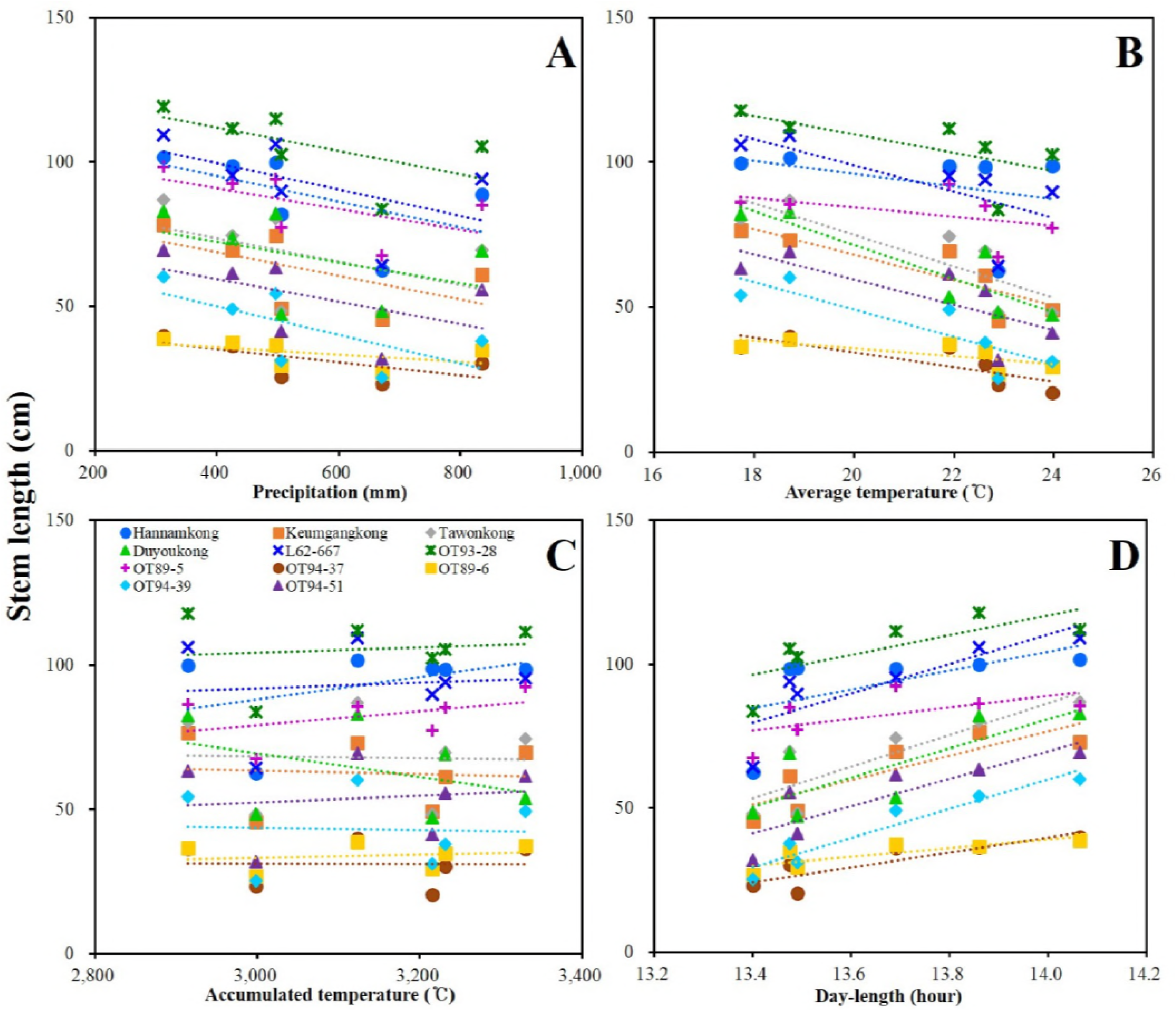
Linear regression analysis between climatic conditions (A: precipitation, B: average temperature, C: accumulated temperature, D: day length) and stem length (STL) of the soybeans grown at six experimental fields in China and the Republic of Korea.

**Fig 7.**
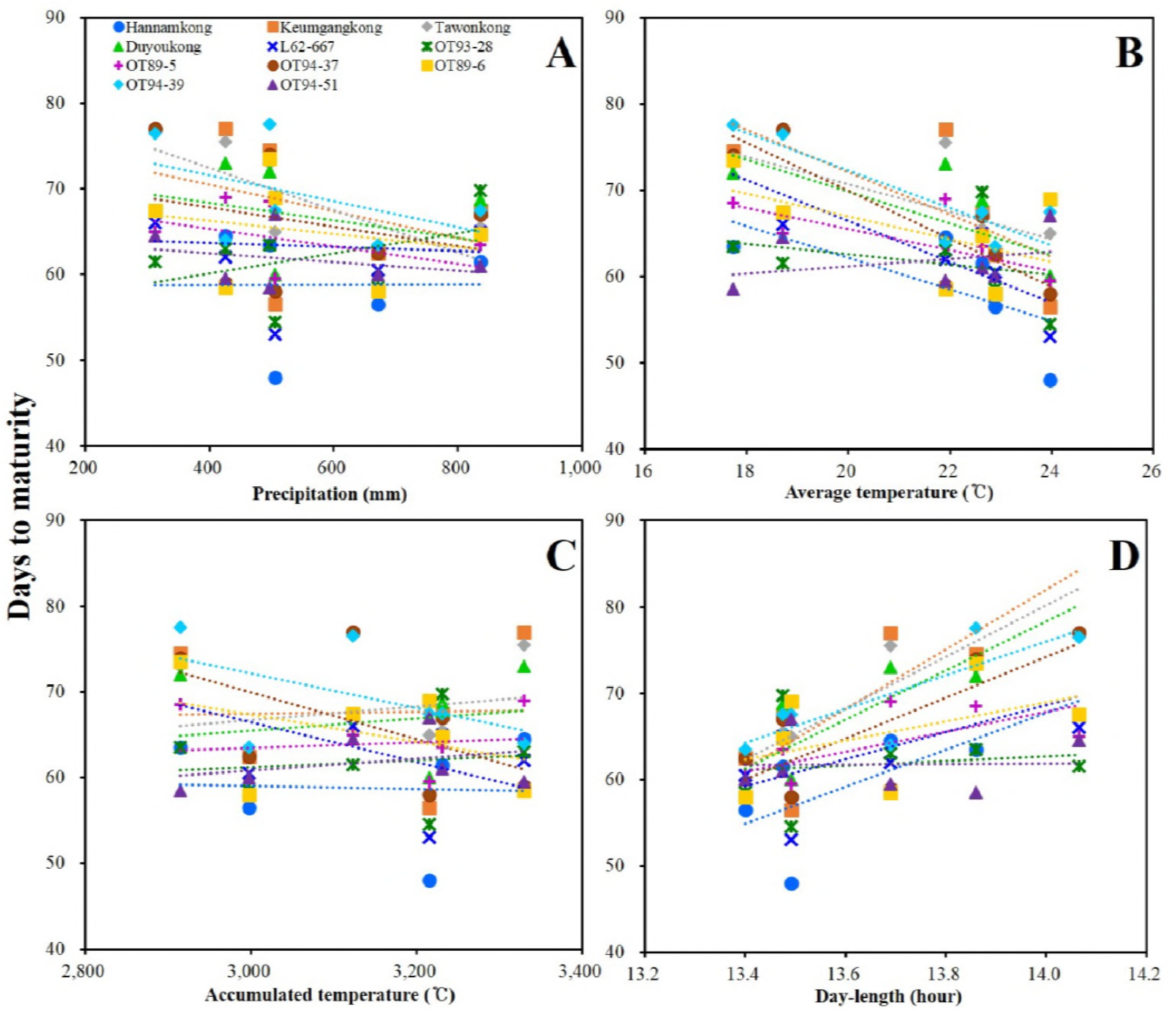
Linear regression analysis between climatic conditions (A: precipitation, B: average temperature, C: accumulated temperature, D: day length) and days to maturity (DTM) of the soybeans grown at six experimental fields in China and the Republic of Korea.

**Fig 8.**
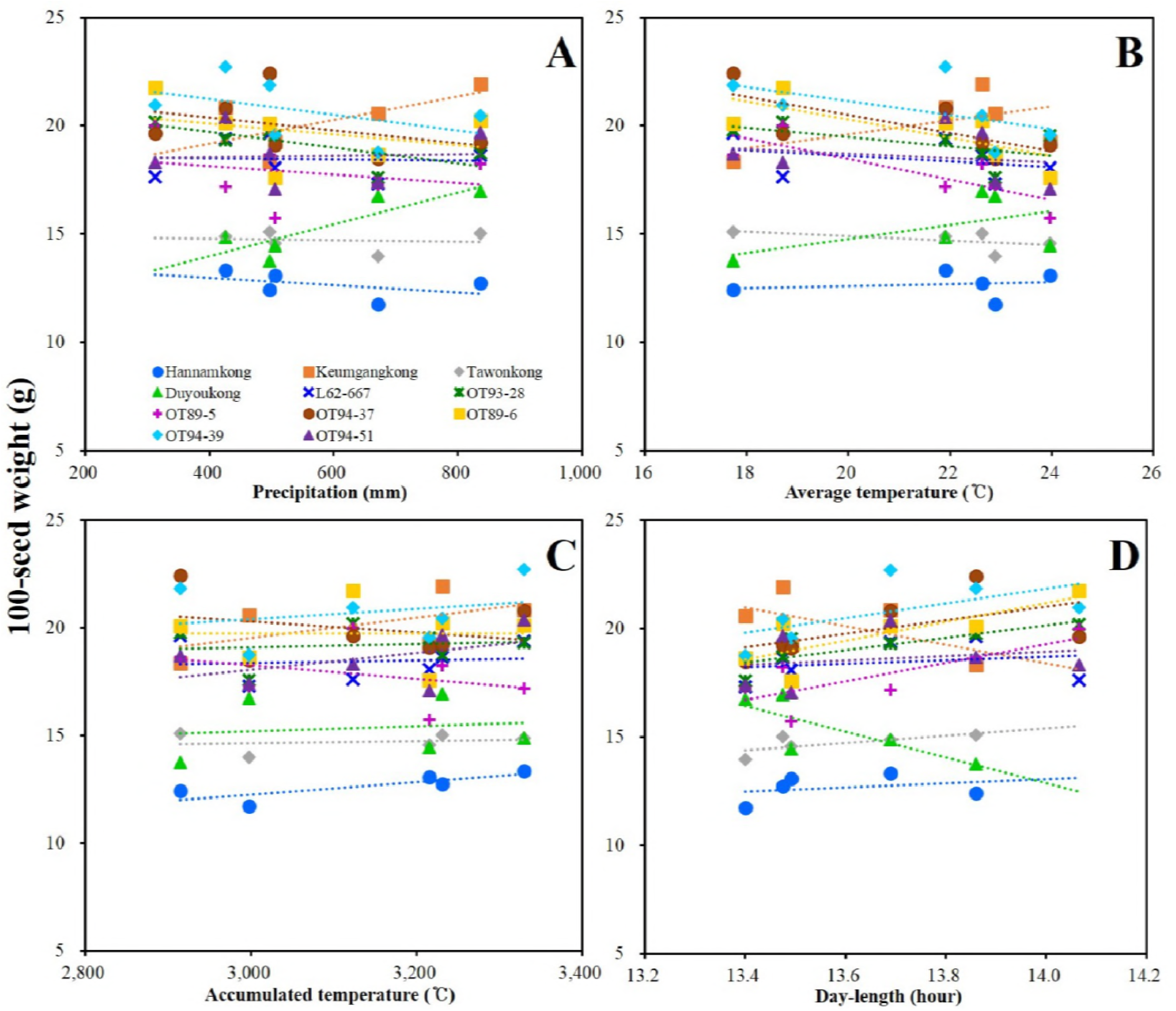
Linear regression analysis between climatic conditions (A: precipitation, B: average temperature, C: accumulated temperature, D: day length) and 100-seed weight (HSW) of the soybeans grown at six experimental fields in China and the Republic of Korea.

**Fig 9.**
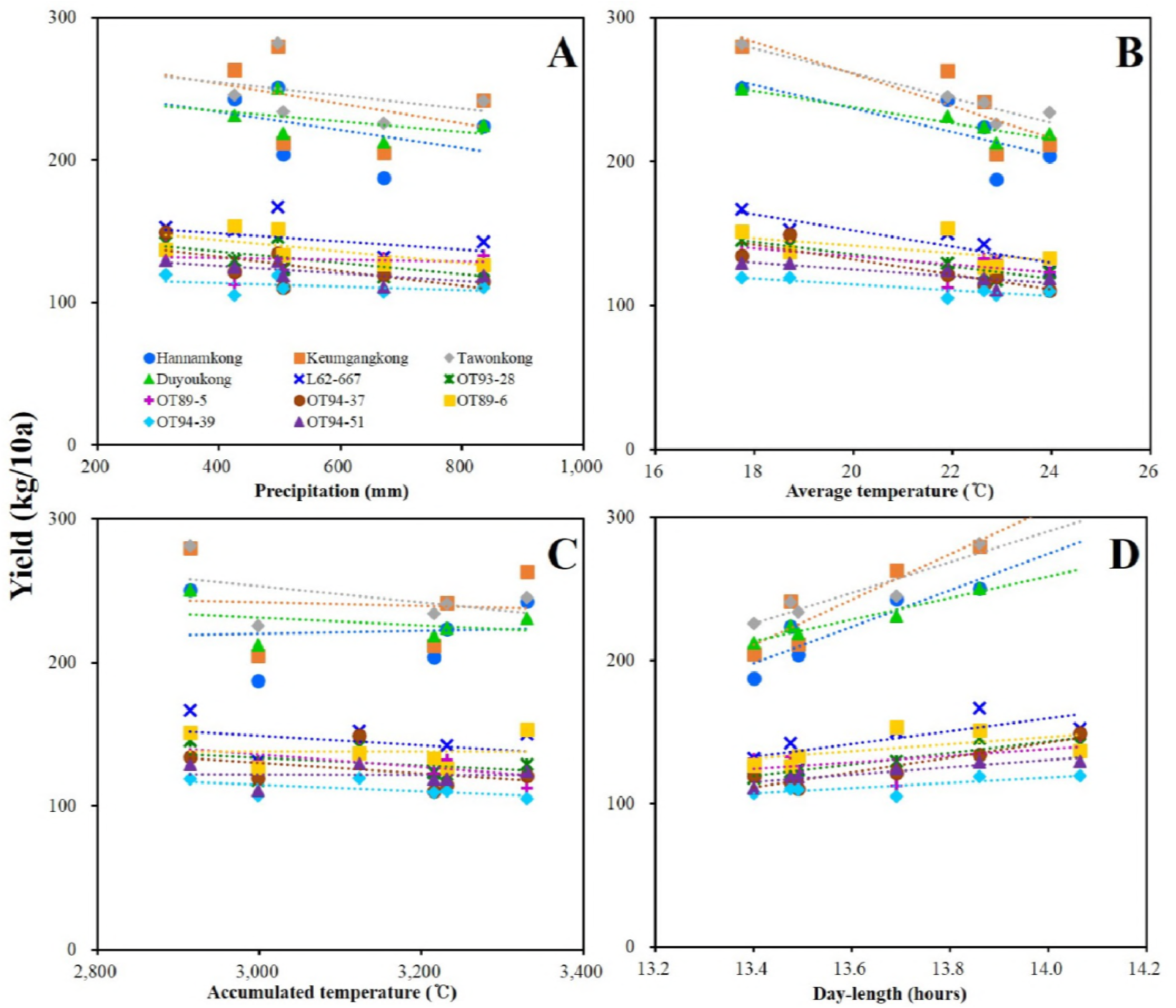
Linear regression analysis between climatic conditions (A: precipitation, B: average temperature, C: accumulated temperature, D: day length) and yield (YLD) of the soybeans grown at six experimental fields in China and the Republic of Korea.

**Table 6.**
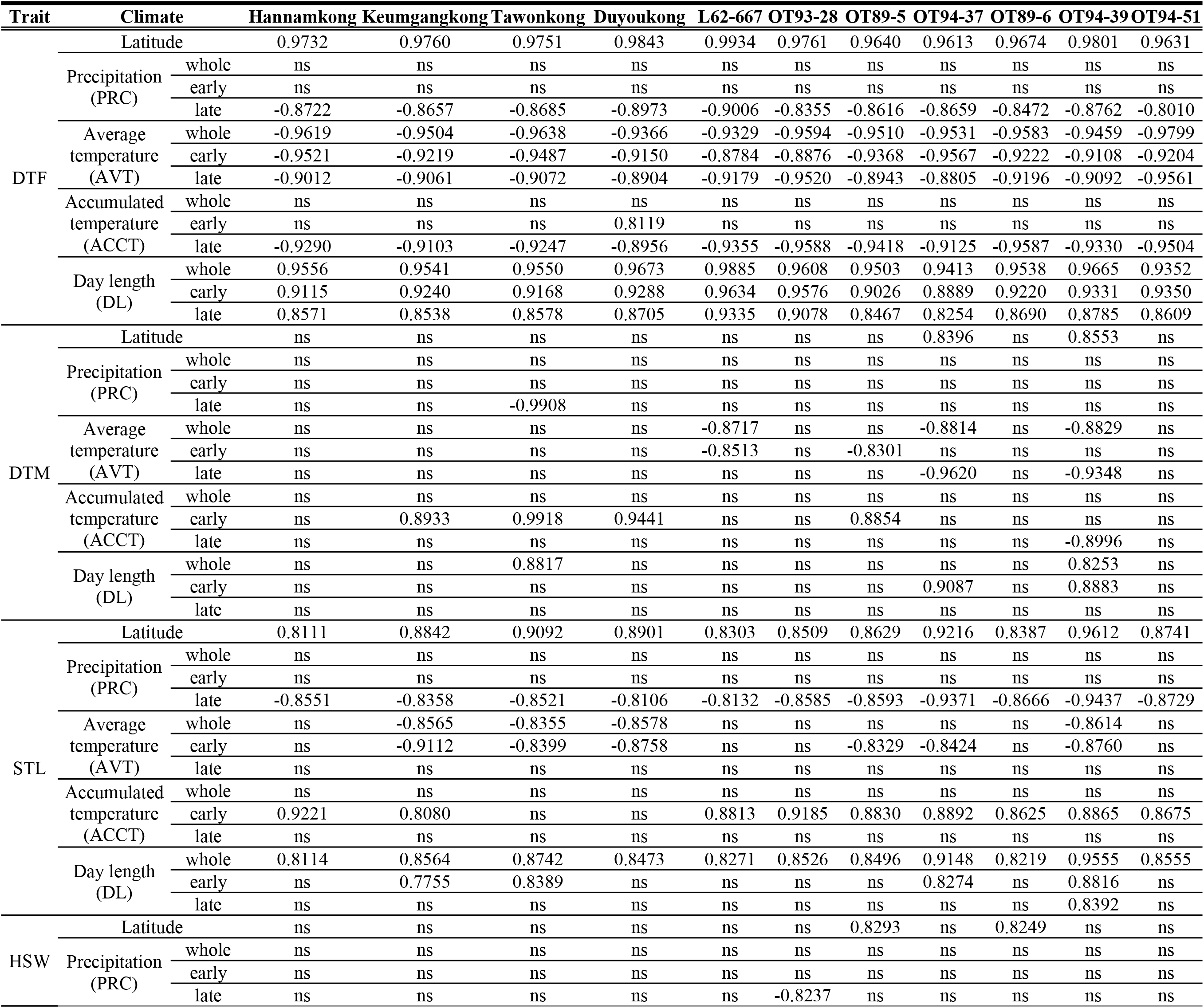
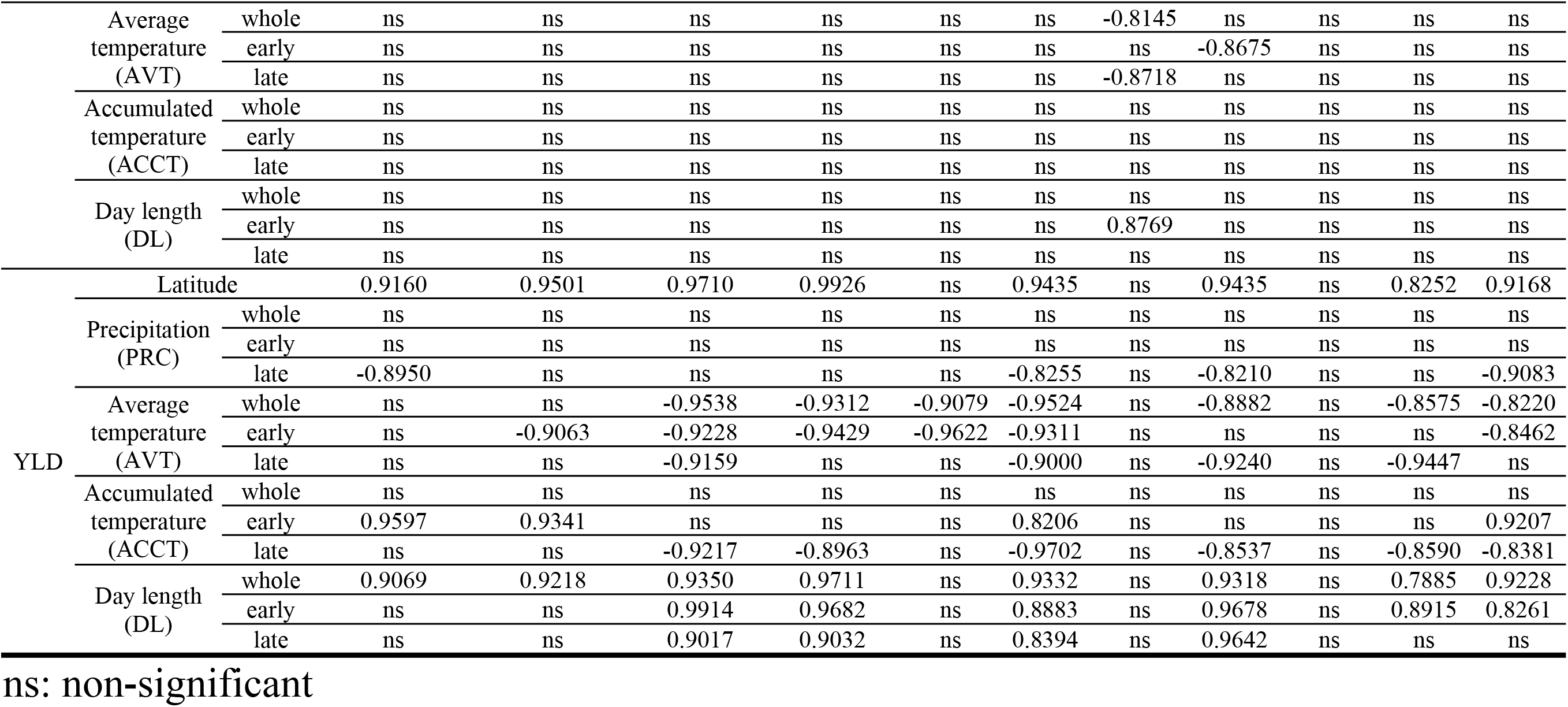
Pearson Correlation Coefficient (PCC) between each climatic condition and days to flowering (DTF), days to maturity (DTM), stem length (STL), 100-seed weight (HSW), and yield (YLD) of the soybeans grown at six experimental sites in China and the Republic of Korea.

### Complementary data

Owing to a temperature of 18°C and 15 h day length, Tawonkong and Hannamkong did not bloom until after the soybeans under other conditions were completely harvested (Fig 10A). Each soybean grown under 23 and 28°C, without considering DL, showed a similar DTF (Fig 10A). The DTF of all soybeans grown at 23 and 28°C under 10 h of DL were almost similar. The DTF of Tawonkong and Hannamkong under 15 h was significantly different from those under 10 h (Fig 10A). Under 15 h DL, as temperature increased, STL of all soybeans lengthened (Fig 10B). Nonetheless, each soybean grown at 23 and 28°C under 10 h DL had a similar STL (Fig 10).

**Fig 10.**
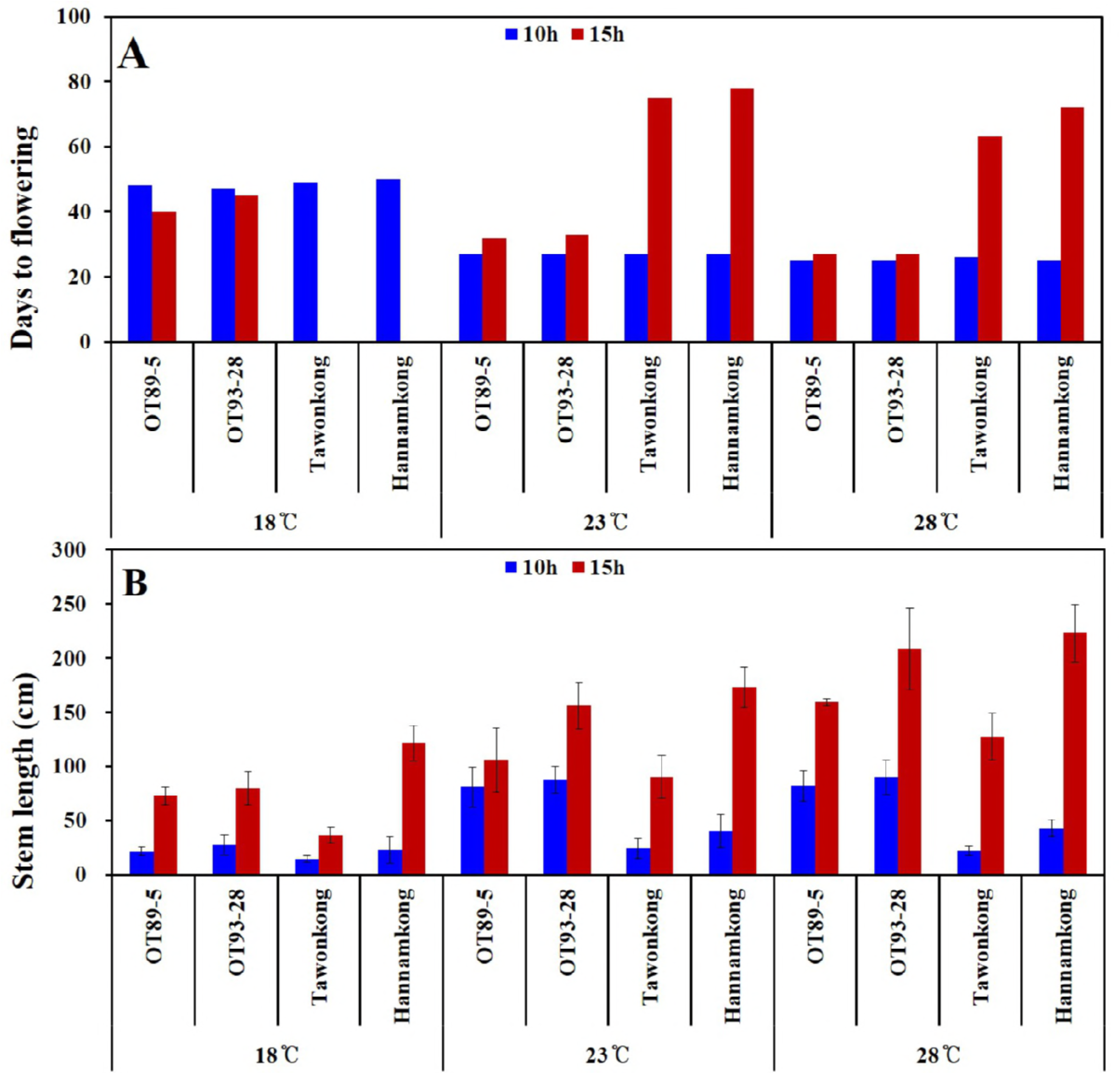
Days to flowering (A) and stem length (B) of the soybeans grown in six growth chambers controlled with six combinations of temperatures (18, 23, and 28°C) and day length (10 and 15 h). LSD (p<0.05) in Fig 10B = 17.325 cm.

## Discussion

### Effect of climatic conditions on flowering and maturity

Climatic conditions including precipitation [8], [10], temperature [13], and day length [5, 6] are highly essential to the agricultural performance of soybean. Thus, understanding the regional climatic characteristic is important for assessing the effects of climatic conditions on the agricultural performances of different soybeans. As expected, the higher the latitude, the higher the DL and the lower the AVT. The latitude showed a negative correlation with PRC (Fig 2, and S2 Table), which had diverse effects on all stages of soybean growth and development.

Fukui and Arai [41] has separately recorded the days from germination to blooming and the days from blooming to ripening with special reference to soybean’s geographical differentiation. This was in an effort to ecologically classify soybeans. Several scientists noted that the days from sowing to R8 (full maturity) have been recorded to classify maturity groups of soybeans based on their response to photoperiod or latitude [42, 43, 44, 45, 46]. However, in previous studies, the days to R1 (DTF) and days to R8 (DTM) were separately calculated to obtain the ratio of R1 to R8 [47], and to estimate the days to R8 as some cultivars did not achieve full maturity before the beginning of the frost period [48]. We recorded DTF and DTM of soybeans separately and examined them in relation to geographical differentiation because we have confirmed via preliminary tests in 2014 and 2015 that some of the cultivars developed in the Republic of Korea were not matured before the beginning of the frost-period in Harbin (Table 4, and Figs 7, 8, and 9).

Xia et al. [49] reported that the E1 allele has a major impact on flowering and maturity. The recessive e1 allele displayed an early flowering time phenotype without reference to the genetic background at other E loci or DL conditions. Moreover, when Zhai et al. [48] classified 180 cultivars into 8 genotypic groups using 4 major E genes (E1 to E4), all soybeans with the e1-nf and e1-as alleles and the E1e2e3e4 genotype were shorter based on time to R1 (under 40 days) than other E1 groups. As mentioned previously, Keumgangkong, Tawonkong and Duyoukong belong to subclade II in the genetic tree constructed by SSR markers associated with DTF. These soybeans having the el-as allele were longer than the other el-as allele genotype soybeans [48, 49] although the DTF was shorter than that of Hannamkong (Table 5, and Figs 4 and 5). Although these results showed that the genetic characteristic of soybean has an effect on DTF, the effect requires further elucidation. As shown in Table 6, although DTF of all the soybeans were highly correlated with all climatic conditions, we could not assume that PRC and ACCT were correlated with DTF as the data for late PRC and late ACCT were measured after R1. The water needs of soybean gradually increase during flowering and through the grain filling stages [8, 11]. For the chamber tests, we confirmed that DTF was negatively and positively correlated with AVT and DL, respectively; the effect of temperature on DTF was not significant when it was over 23°C (Fig 10). However, the average of the early AVT at the six fields was 22.6°C (Table 3). Based on these results, we concluded that although the DTF of soybean grown in Northeast Asia is affected by its genetic characteristic, AVT, and DL, the effect of DL is more significant than the others.

Between the subclades of the tree structure shown in Fig 4B, and the data for the DTM (R1 to R8) shown in Table 4, no correlation was found. In addition, as can be seen in Fig 7 and Table 6, the DTM was not significantly correlated with any climatic conditions. As shown in Table 4, Hannamkong showed the longest DTF and the shortest DTM. Moreover, the average of the DTM of soybeans including OT94-37, OT89-6, and OT94-39 was the longest, as is seen in subclade I. In the tree associated with DTF, these soybeans were classified into subclades III and IV, which had the shortest average of DTF (Table 4, and Fig 4A and 4B). These results showed that the internal and external influences on DTM of soybean require further elucidation.

### Effect of climatic conditions on stem length

Stem termination has an effect on plant height, flowering period, node production, maturity, water-use efficiency, and yield of soybean [23, 50, 51]. Genotyping using the determinate stem (Dt1) allele plays an essential role in analysis of genetic characteristic associated with the stem length and maturity of soybean. The soybeans with a dominant Dt1 allele (except Tawonkong) and classified into subclades III and VI of the tree associated with STL had the longest average of STL (Tables 4 and 5, and Fig 4C), as in previous reports [23, 52]. These results imply that the genetic characteristic has a large effect on the STL of soybean. The results of our field tests showed that STL has a positive correlation with DL, while AVT did not. Our chamber test confirmed that STL of soybean is positively affected by temperature and DL, results seen in previous studies [53, 54]. However, the result that the correlations between STL and AVT in the field and chamber tests were different was caused by the characteristic of regional climates where latitude showed a negative correlation with AVT (Fig 2, S1 Table). Therefore, these results showed that STL of soybean is more affected by its genetic characteristic and DL than AVT, as with DTF.

### Effect of climatic conditions on yield components

In the genetic tree associated with HSW, 3 subclades with a similar HSW were formed, as also seen in the tree associated with YLD (Fig 4D and 4E). Many researchers have reported that HSW and YLD of soybean have the same responses to climatic conditions such as latitude, PRC, temperature, and DL [32, 55, 56, 57]. Nevertheless, HSW showed no correlation with any climatic conditions although the YLD of the soybeans, except L62-667, OT89-5, and OT89-6, showed positive correlations with latitude and DL. As shown in Fig 9, the soybeans were distinctly divided into two groups according to YLD, without reference to climatic conditions. This indicates that YLD of soybean is mostly affected by its genetic characteristic. The soybeans that had a relatively longer DTF (over 55 days) showed higher correlations with DTF than others which had relatively short DTFs (under 47 days). In addition, our study showed that YLD has a positive correlation with DTF (Fig 3). These results indicate that DTF has a large effect on YLD. Nonetheless, Yamada et al. [58] reported two different results related to DTF and YLD of soybeans grown in the southern and northern areas of Japan. In the northern area, YLD of the soybeans that had a relatively shorter DTF was higher than that of the others that had a relatively longer DTF. The second result, however, was similar to our findings. Therefore, for these results, we concluded that climatic conditions selectively affects YLD based on the length of DTF; the response of soybean YLD to climatic conditions is cultivar-specific, and more specifically based on genetic characteristic.

The DTF and STL of soybeans grown in Northeast Asia, China and the Republic of Korea, are affected by DL and AVT as well as genetic characteristic. DL, however, had a greater effect on DTF and STL than it had on other traits such as DTM, HSW, and YLD. Although their cultivar-specific characteristic greatly contributes to YLD of soybeans, the YLD of the soybeans grown in Northeast Asia selectively responds to the latitude and DL based on the length of DTF.

This study assessed the independent and interactive effects of climatic conditions on the agricultural performance of soybeans grown at six different latitudes in China and the Republic of Korea, in an effort to understand the phenological development of soybeans in these two countries. The study findings contribute to our understanding of the consequent effects of breeding and selection of soybeans under climate change in the two countries.

## Supporting information

S1 Table. Correlation coefficient values and p-values for agricultural performance of the soybeans grown at six experimental fields in 2016-2017.

S2 Table. Correlation coefficient values and p-values for climatic conditions at six experimental fields investigated in 2016-2017.

## Acknowledgments

This study was supported by the Rural Development Administration (RDA, Republic of Korea) grant PJ012657032018.

## Author Contributions

Conceptualization: Myoung Ryoul Park.

Data curation: Myoung Ryoul Park.

Formal analysis: Myoung Ryoul Park.

Investigation: Myoung Ryoul Park, Chunmei Cai, Man-Soo Choi, Soo-Kwon Park, Chang-Hwan Park, Jung Kyung Moon.

Methodology: Myoung Ryoul Park, Chunmei Cai.

Software: Myoung Ryoul Park.

Supervision: Myoung Ryoul Park, Chunmei Cai.

Validation: Min-Jung Seo, Hong-Tae Yun.

Visualization: Myoung Ryoul Park.

Writing - original draft: Myoung Ryoul Park.

Writing - review & editing: Myoung Ryoul Park, Chunmei Cai, Min-Jung Seo, Man-Soo Choi, Soo-Kwon Park, Hong-Tae Yun, Chang-Hwan Park, Jung Kyung Moon.

